# Mechanism of SARS-CoV-2 Nucleocapsid Protein Phosphorylation-induced Functional Switch

**DOI:** 10.1101/2025.09.12.675721

**Authors:** Megan S. Sullivan, Michael Morse, Kaylee Grabarkewitz, Dina Bayachou, Ioulia Rouzina, Vicki Wysocki, Mark C. Williams, Karin Musier-Forsyth

## Abstract

The SARS-CoV-2 nucleocapsid protein (Np) is essential for viral RNA replication and genomic RNA packaging. Phosphorylation of Np within its central Ser-Arg-rich (SRR) linker is proposed to modulate these functions. To gain mechanistic insights into these distinct roles, we performed in vitro biophysical and biochemical studies using recombinantly expressed ancestral Np and phosphomimetic SRR variants. Limited-proteolysis showed minor cleavage differences between wild-type (WT) and phosphomimetic Np, but no major structure or stability changes in the N- and C-terminal domains were observed by circular dichroism spectroscopy and differential scanning fluorimetry, respectively. Mass photometry (MP) revealed that WT Np dimerized more readily than phosphomimetic variants. Crosslinking-MP showed WT Np formed discrete complexes on viral 5ʹ UTR stem-loop (SL) 5 RNA, whereas phosphomimetic Np assembled preferentially on SL1-4. WT Np bound non-specifically to all RNAs tested primarily via hydrophobic interactions, whereas phosphomimetic Np showed selectivity for SARS-CoV-2-derived RNAs. WT Np also compacted and irreversibly bound single-stranded DNA; this activity was significantly reduced by phosphorylation. These mechanistic insights support a model where phosphorylated Np functions in RNA replication and chaperoning, while non-phosphorylated Np facilitates genomic RNA packaging. The findings also help to explain infectivity differences and clinical outcomes associated with SRR linker variants.

## INTRODUCTION

Severe Acute Respiratory Syndrome Coronavirus 2 (SARS-CoV-2) is a betacoronavirus and the causative agent of the COVID-19 pandemic. Coronaviruses are enveloped viruses with virions assembled by four structural proteins: spike (S), envelope (E), membrane (M), and nucleocapsid (N). They encode large (∼30 kb) positive-sense, single-stranded RNA (+ssRNA) genomes, and the viral N protein (Np) is responsible for packaging this full-length RNA into progeny virions (1,2).

The ∼45.7 kDa Np consists of an N-terminal intrinsically-disordered region (IDR), a structured N-terminal domain (NTD), a central IDR linker region, a structured C-terminal domain (CTD), and a C-terminal IDR (3–8). The NTD is primarily responsible for RNA binding, whereas the CTD is responsible for protein dimerization and RNA binding. The linker IDR possesses a conserved Ser-Arg rich (SRR) hyperphosphorylated region and a leucine helix (LH) domain (9). Upon viral entry into the host cell, the +ssRNA is directly translated into polyprotein1a or 1ab (10,11). These polyproteins are autoproteolytically cleaved into non-structural proteins (nsps) that enable formation of double-membraned vesicles (DMVs), a viral organelle that houses the viral replication-transcription complex (RTC), also made of nsps (10,12–14).

Full-length gRNA replication and synthesis of all accessory and structural protein mRNA transcripts occurs at the RTC through a process of discontinuous transcription (15,16). Briefly, the viral RNA-dependent RNA polymerase (RdRp) begins synthesis of a negative-sense daughter strand at the 3’ end of the genome. A conserved six-nucleotide (nt) sequence, the body transcription regulatory sequence (B-TRS) is present between each gene-coding region. The same six-nt sequence occurs at positions 70-75 in the viral RNA 5’ untranslated region (5’UTR) and is referred to as the leader TRS (L-TRS). Upon arrival at the end of each gene, the RdRp will either continue transcription through the B-TRS, or “jump” ∼20,000 nt to the L-TRS where it then adds the first 70 nt of the 5’UTR to the daughter strand. The newly-transcribed RNAs are referred to as negative-sense subgenomic mRNAs (-sgmRNAs). The -sgmRNAs are used as templates by the RTC to produce +sgmRNAs, which are exported through the Arg-lined DMV pore to be translated (17). Although the general mechanism of discontinuous transcription is widely accepted, the regulation of RdRp activity remains poorly understood (15,16). The viral Np has been proposed to facilitate this process by acting as an RNA chaperone, independently of its role in genome packaging.

RNA chaperone proteins aid in RNA structure remodeling through destabilization of secondary structure (entropy transfer model), whereas RNA annealers aid in RNA base pairing through molecular crowding and/or RNA secondary structure remodeling that exposes surfaces for base pairing to then take place (18,19). SARS-CoV, transmissible gastroenteritis virus (TGEV), and murine hepatitis virus (MHV) Nps have all demonstrated RNA chaperone and RNA annealing activity (20–23). Geidroc and coworkers characterized the thermodynamics of SARS-CoV and MHV NTD domain binding to TRS-containing helices. The MHV NTD specifically and robustly melted 15-mer RNA duplexes containing the leader TRS on one strand and the complementary sequence (cTRS) on the second strand, and the SARS-CoV NTD completely melted a 10-mer RNA TRS-cTRS duplex (22,23). Using an HIV-1 derived strand-transfer assay, Zuniga *et al.* demonstrated both SARS-CoV Np and TGEV Np enhance the self-cleavage activity of the avocado sunblotch viroid ribozyme and promote the annealing of TRS-cTRS RNA oligonucleotides (20). They also demonstrated that TGEV Np can facilitate strand transfer ∼3-fold above a glutathione *S*-transferase (GST) background control (21). Full-length SARS-CoV-2 Np was shown to anneal HIV-1 derived nucleic acids to a similar extent as HIV-1 NCp9, a well-characterized viral nucleic acid chaperone protein (24,25). The role of SRR phosphorylation in Np chaperone function has not been directly tested.

Studies of MHV and SARS-CoV demonstrated that virions contain exclusively non-phosphorylated Np, but Np phosphorylation is required for RTC synthesis of the longest sgmRNA transcripts and for maintaining high infectivity (26–28). The SRR of SARS-CoV-2 Np has also been shown to associate with the N-terminal ubiquitin-like domain 1 (Ubl1) of viral nsp3, a component of the DMV pore. Changing the phosphorylation state of the SRR (via either S to A or S to D mutations) disrupted this interaction and decreased viral replication (7,29–31). More recent *in vitro* studies demonstrated phosphorylation-dependent differences in the liquid-liquid phase-separation (LLPS) behavior of SARS-CoV-2 Np. In the presence of RNA, phosphorylated Np formed liquid-like droplets, whereas non-phosphorylated Np formed more gel-like, rigid, branched structures (32–37). Np condensation with RNA required the central IDR; both a phosphoablative-SRR construct and an SRR-deletion construct expressed in mammalian cells formed viscous granules with RNA in contrast to WT or phosphomimetic Np, which formed condensates that remained liquid-like. Thus, SRR phosphorylation status is critical for modulating Np-RNA LLPS behavior (37). Taken together, these data suggest that dynamic phosphorylation of SARS-CoV-2 Np is a means by which its two major functions, genomic RNA packaging and facilitation of -sgmRNA synthesis, are modulated (34,36,38–40). However, the molecular mechanism of this Np functional switch is still unclear.

In this work, we tested the mechanism by which SRR phosphorylation impacts SARS-CoV-2 Np function by probing the effect of phosphorylation on Np protein structure and stability, as well as nucleic acid interaction properties. Our data suggest that phosphorylation of Np changes the RNA binding mode from a robust condensing agent (non-phosphorylated) into a dynamic chaperone protein (phosphorylated). This work has implications for better understanding the role of Np in coronavirus infection and supports the growing body of work showing Np phosphorylation is a critical modification for viral replication.

## MATERIAL AND METHODS

### RNA preparation

#### a. Plasmid constructs and in vitro transcription template preparation

A pIDTsmart plasmid vector encoding the first 300 nucleotides of the SARS-CoV-2 genome (5ʹ UTR) with an upstream T7 promoter sequence, a hammerhead ribozyme for correct 5ʹ end generation, and a 3ʹ end FokI digestion site was commercially purchased from IDT (41). To prepare DNA template for in vitro transcription, plasmid was purified using a MaxiPrep (Qiagen) purification kit, according to the manufacturer’s protocol, and linearized via FokI digestion (New England Biolabs). Linearized template DNA was precipitated with isopropanol, resuspended in MilliQ-H_2_O and stored at −20°C.

Linearized template DNA was used for PCR amplification of SL1-4 and SL5 DNA templates with Phusion DNA polymerase (New England Biolabs) and primers purchased from IDT. For both constructs, the reverse primers included a single 5ʹ-end 2ʹ-O-methyl modification to reduce T7 3ʹ runoff transcription activity (42). SL1-4 primers amplified 1-148 nt and SL5 primers amplified 149-300 nt of the viral 5ʹ UTR. The forward SL5 primer also contained the T7 promoter sequence. PCR products were purified using PCR clean-up kit (Qiagen) and stored at −20°C. HIV-1 TARpolyA and WT Psi DNA constructs were previously described (43).

#### b. In vitro transcription and purification of viral 5’UTR RNAs

To make SL1-5 RNA, 100 µg of FokI-linearized template plasmid DNA was used in a 1 mL transcription reaction containing 80 mM HEPES pH 8.0, 30 mM Mg(OAc)_2_, 10 mM dithiothreitol (DTT), 5 mM spermidine, 0.01% Triton X-100, 4 mM each of ATP, UTP, and CTP, 8 mM GTP, 5U of yeast inorganic pyrophosphatase (New England Biolabs or in-house purified), and recombinant, in-house purified P266L T7 RNA polymerase; enzyme purifications were performed as described (44,45). SL1-4 and SL5 DNA templates were similarly used to prepare these RNAs by in vitro transcription except using ∼15 μg of PCR-amplified DNA per 1 mL transcription reaction. Plasmid pET29b-IPP1-His encoding yeast inorganic pyrophosphatase was a gift from Sebastian Maerkl & Takuya Ueda (Addgene plasmid # 124137; RRID: Addgene_124137), and plasmid pBH161 encoding P266L T7 RNA polymerase was a gift from the Gopalan lab at Ohio State University. After incubation at 37°C for 4 h, reactions were quenched with addition of 50 mM EDTA, pH 8.0 (Invitrogen, UltraPure^TM^). The RNA was then either gel-purified and eluted via the crush and soak method, FPLC purified via size-exclusion chromatography, or HPLC purified via ion-pair reverse-phase chromatography (46–49). For all liquid chromatography methods, RNA was extracted with acidic phenol:chloroform, ethanol precipitated, and resuspended in MilliQ-H_2_O before column application.

Size-exclusion chromatography was performed using buffer containing 10 mM Tris-HCl pH 6.5, 50 mM NaCl on a HiPrep 16/60 Sephacryl S 400 HR column (Cytiva) using an ATKA Pure FPLC (Cytiva) instrument (47). RNA was eluted over 1.5 column volumes, and the fractions containing RNA were identified using 6% urea-PAGE, combined, and then ethanol precipitated. The pellet was resuspended in MilliQ-H_2_O, the concentration determined via A_260_, and product stored at −20°C. The RNA extinction coefficients (χ_260_) used are listed in Table 1.

**Table 1.** Molecular weights and extinction coefficients of proteins and RNAs.

| Protein | $\epsilon_{280}$ | Molecular weight | RNA | $\epsilon_{260}$ | Molecular weight |
| --- | --- | --- | --- | --- | --- |
| WT Np | 43,890 M<sup>-1</sup>·cm<sup>-1</sup> | 45,683 Da | SL1-5 | 2,757,968 M<sup>-1</sup>·cm<sup>-1</sup><sub>1</sub> | 102,000 Da |
| 3D1 Np | 43,890 M<sup>-1</sup>·cm<sup>-1</sup> | 45,767 Da | SL1-4 | 1,346,873 M<sup>-1</sup>·cm<sup>-1</sup><sub>1</sub> | 50,660 Da |
| 3D2 Np | 43,890 M<sup>-1</sup>·cm<sup>-1</sup> | 45,753 Da | SL5 | 1,374,908 M<sup>-1</sup>·cm<sup>-1</sup><sub>1</sub> | 51,660 Da |
| 6D Np | 43,890 M<sup>-1</sup>·cm<sup>-1</sup> | 45,837 Da | TARpolyA | 945,038 M<sup>-1</sup>·cm<sup>-1</sup> | 36,720 Da |
| HIV-1 Gag | 65,890 M<sup>-1</sup>·cm<sup>-1</sup> | 58,051 Da | WT Psi | 963,728 M<sup>-1</sup>·cm<sup>-1</sup> | 36,040 Da |

Ion-paired reverse phase HPLC purification of RNA was performed on an Agilent 1260 HPLC instrument using a Poroshell 120 C18 (Agilent) column equilibrated with 2% acetonitrile, 0.1 M triethylamine-acetate (TEAA) pH 7.0. Before injection, RNA samples were heat denatured at 95°C for 2 min and filtered through a 0.2 µm polyethersulfone (PES) syringe filter (Whatman). For each injection, ∼5 µL of sample was run and RNA was eluted using a linear acetonitrile gradient from 5%-12% (48,49). Fractions to collect were identified using A_260_ detection. After RNA elution, the column was washed with a linear gradient of acetonitrile in MilliQ-H_2_O from 20%-50%, then re-equilibrated with a buffer containing 2% acetonitrile, 0.1 M TEAA pH 7.0, before the next injection. All reagents used were HPLC grade obtained from Sigma. Collected fractions were analyzed using 6% urea-PAGE, and fractions containing RNA were combined, butanol extracted to reduce the volume, and ethanol precipitated. The pellet was resuspended in MilliQ-H_2_O, the concentration determined via A_260_, and product stored at −20°C.

### Protein preparation

A pET28⍺ vector encoding SARS-CoV-2 ancestral WT-Np containing an N-terminal TEV-cleavable 10x-His tag and kanamycin resistance cassette was obtained as a gift from Stefan Sarafinos (Emory School of Medicine). Phosphomimetic mutants (3D1 Np: S176D, S180D, and S184D; 3D2 Np: T198D, S202D, and S206D; 6D Np: S176D, S180D, S184D, T198D, S202D, and S206D) were generated using site-directed, ligase-independent mutagenesis (SLIM) with the WT-Np plasmid as a template and primers from Integrated DNA Technologies (IDT) (Table S1) (50,51). The sequences were confirmed using Sanger sequencing. *E. coli* DH5⍺ and Rosetta (DE3) competent cells were transformed with WT and mutant plasmids for storage and protein purification, respectively.

Recombinant WT HIV-1 Gag was purified as described (52). Recombinant SARS-CoV-2 WT Np or phosphomimetic proteins were expressed in *E. coli* Rosetta (DE3) cells using ZYM-5052 autoinduction media and antibiotic selection with both chloramphenicol and kanamycin (53). Cell pellets were resuspended in a buffer containing 50 mM HEPES pH 7.5, 1 M NaCl, 5 mM β-mercaptoethanol (βME), 5% (v/v) glycerol, and 0.5% (v/v) Triton X-100 with approximately 100 mg of lysozyme and 1 tablet of Roche cOmplete, EDTA-free protease inhibitor per 50 mL buffer. Cells were lysed using sonication, followed by addition of 1 µL of DNase I (New England Biolabs) per 1 mL of lysate and incubation with stirring at 4°C for 30-60 min. Total lysate was then centrifuged at 15,000 xg at 4°C for 30 min. The supernatant was used for native purifications of 3D1 and 6D proteins, and the insoluble pellet was used for denaturing purifications of 3D2 and WT Np. The latter had very low yield and considerable nucleic acid contamination with the native purification method, so a denaturing purification protocol adapted from Dieci *et al* (2000) and Cabrita and Bottomley (2004) was developed (54,55).

#### a. Native purification method

Total lysate supernatant was passed through a 0.45 µm filter followed by two ammonium sulfate precipitations. First, 5 M (NH_4_)_2_SO_4_ was added to 10% saturation to precipitate contaminating proteins. Next, to precipitate Np from the resulting supernatant, (NH_4_)_2_SO_4_ was added to 33% final saturation. The solution was centrifuged at 10,000 xg, at 4°C for 20 min, with a slow deceleration to prevent pellet disintegration. The pellet was resuspended in 50 mL of Ni^2+^-affinity column equilibration buffer (50 mM HEPES pH 7.5, 0.5 M NaCl, 15 mM imidazole, 5 mM βME, 5% (v/v) glycerol) then centrifuged again to remove any remaining precipitate. The supernatant was loaded onto a gravity column at 4°C containing Pierce^TM^ High-capacity Ni-IMAC resin, EDTA compatible (Thermo Scientific). Resin was washed with 2 column volumes of the equilibration buffer containing 50 mM imidazole, and protein was eluted using a step-wise gradient of 150 mM, 210 mM, 300 mM and 500 mM imidazole. Fractions were analyzed using 10% SDS-PAGE, and those containing Np were combined. To cleave the His-tag, 1 mg of in-house purified His_6_-TEV protease was added to the combined fractions and dialyzed using a Slide-A-Lyzer 2.5 kDa MWCO cassette (Thermo Scientific) with gentle stirring at 4°C overnight in 50 mM HEPES pH 7.5, 300 mM NaCl, 5 mM βME, 5% (v/v) glycerol. TEV cleavage was confirmed using 10% SDS-PAGE before proceeding with a second Ni-IMAC column purification to remove the cleaved His-tag, any remaining uncleaved protein, and the TEV protease. The flow through and low-imidazole (20 mM) wash were analyzed via 10% SDS-PAGE to confirm that only uncleaved N-protein was present. If protein purity was less than 95%, the combined fractions were subjected to gravity cation exchange chromatography using Fractogel EMD SO_3_^-^ (M) resin (Millipore). After sample loading, the column was first washed with 1 column volume of equilibration buffer described above but containing 300 mM NaCl, followed by 1 column volume with the same buffer containing 0.5 M NaCl. Protein was eluted using a stepwise gradient of 0.75 M, 0.85 M, and 1 M NaCl. Fractions were analyzed via 10% SDS-PAGE, and those containing protein were combined. The purified protein was buffer exchanged into storage buffer (20 mM HEPES pH 7.5, 0.5 M NaCl, 5% (v/v) glycerol, 1 mM DTT) and concentrated using a VivaSpin-20, 30 kDa MWCO centrifugal concentrator (Cytiva). Protein concentrations were determined using A_280_, aliquoted, and stored at −80°C. The concentration was also verified via SDS-PAGE using a BSA mass standard, and adjustments to the concentration were made if necessary through ImageJ densitometry gel quantification. Protein extinction coefficients and molecular weights used for quantification are listed in Table 1 and were obtained using the ExPASy ProtParam online tool (56).

#### b. Denaturing purification method

The cell lysate insoluble pellet was resuspended in 20 mL of inclusion body lysis buffer consisting of 20 mM HEPES pH 8.0, 0.3 M NaCl, 6 M guanidine-HCl, 5 mM βME, and 1 tablet of Roche cOmplete EDTA-free protease inhibitor. The resuspended lysate was stirred for 1 h at room temperature (22-23°C) and then clarified via centrifugation at 15,000 xg at 4°C for 30 min. The clarified lysate was subjected to Ni^2+^-affinity chromatography using Pierce^TM^ high-capacity Ni-IMAC EDTA-compatible resin at room temperature to prevent urea crystallization during wash steps. The column was equilibrated in inclusion body lysis buffer, and clarified lysate was run through the column. The flow through was collected and loaded to the column a second time to ensure maximum protein binding. The column was washed with a series of urea washes (3 column volumes each of 8 M, 7 M, 6 M, 5 M, and 4 M urea), all containing 20 mM HEPES pH 8.0, 300 mM NaCl, 1% (w/v) CHAPS (GoldBio), 5 mM βME and 10 mM imidazole. Denatured Np was eluted using elution buffer (20 mM HEPES pH 8.0, 300 mM NaCl, 1% (w/v) CHAPS, 5 mM βME, 500 mM imidazole, 4 M urea). Fractions were analyzed via 10% SDS-PAGE, and those containing Np were combined and subjected to gentle dialysis (30 kDa MWCO Slide-a-lyzer dialysis cassette) against 2 L of a series of refolding dialysis buffers over the course of 3 total days (see Table 2). Once dialysis into buffer 6 was complete, Np was removed from the dialysis cassette and any precipitated/aggregated protein was pelleted by a short centrifugation step (4198 xg, 4°C, 1 minute) in a swinging bucket centrifuge. Supernatant was removed, and approximately 1 mg of His_6_-TEV was added followed by overnight dialysis using a 2.5 kDa MWCO Slide-a-lyzer cassette against buffer 7 (see Table 2). TEV cleavage was verified via 10% SDS-PAGE before proceeding with a second Ni-IMAC column to remove the cleaved His-tag, uncleaved protein, and TEV protease. Protein was pooled after fractions were verified using 10% SDS-PAGE and then buffer exchanged into storage buffer using a VivaSpin-20, 30 kDa MWCO centrifugal concentrator. Protein concentration was determined via A_280_, aliquoted, and stored as described above.

**Table 2.** Denaturing purification refolding dialysis buffers.

| Buffer number | Contents (2 L total volume each) |
| --- | --- |
| 1<br>Day 1 | 20 mM HEPES pH 8.0, 0.5 M NaCl, 5% (v/v) glycerol, 0.1 mM TCEP, 4 M urea, 50 mM imidazole, 0.5% (w/v) CHAPS, 0.1 M glutamate + arginine pH 8.0 |
| 2<br>Day 1 | 20 mM HEPES pH 8.0, 0.5 M NaCl, 5% (v/v) glycerol, 0.1 mM TCEP, 3 M urea, 37.5 mM imidazole, 0.5% (w/v) CHAPS, 0.1 M glutamate + arginine pH 8.0 |
| 3<br>Day 1 | 20 mM HEPES pH 8.0, 0.5 M NaCl, 5% (v/v) glycerol, 0.1 mM TCEP, 2 M urea, 25 mM imidazole, 0.33% (w/v) CHAPS, 0.1 M glutamate + arginine pH 8.0 |
| 4<br>Day 2 | 20 mM HEPES pH 8.0, 0.5 M NaCl, 5% (v/v) glycerol, 0.1 mM TCEP, 1 M urea, 17.5 mM imidazole, 0.167% (w/v) CHAPS, 88 mM glutamate + arginine pH 8.0 |
| 5<br>Day 2 | 20 mM HEPES pH 8.0, 0.5 M NaCl, 5% (v/v) glycerol, 0.1 mM TCEP, 0.75 M urea, 13 mM imidazole, 0.125% (w/v) CHAPS, 66 mM glutamate + arginine pH 8.0 |
| 6<br>Day 2 | 20 mM HEPES pH 8.0, 0.5 M NaCl, 5% (v/v) glycerol, 0.1 mM TCEP, 0.5 M urea, 5 mM imidazole, 0.083% (w/v) CHAPS, 45 mM glutamate + arginine pH 8.0 |
| 7 (TEV<br>cleavage) | 20 mM HEPES pH 8.0, 0.5 M NaCl, 5% (v/v) glycerol, 0.1 mM TCEP, 0.25 M urea, 5 mM imidazole, 0.021% (w/v) CHAPS, 23 mM glutamate + arginine pH 8.0 |

### Fluorescence anisotropy (FA) salt-titration assays

Viral 5’UTR RNAs listed in Table 1 were fluorescently labeled on the 3’ end with fluorescein-5-thiosemicarbazide (FTSC) (Invitrogen) as previously described, except after the labeling reaction, the solution was run through a G25 desalting column (Thermo Scientific) (43). The concentration of fluorophore was calculated using A_495_, and concentration of the labeled RNA was quantified by adjusting the A_260_ with the FTSC stock correction factor (A_260_/A_495_) as described (43). NaCl titration assays were performed as described using a BioTek Cytation 5 plate reader outfitted with a fluorescence polarization filter cube (510 nm dichroic mirror; 485 nm excitation, 528 nm emission, each with 20 nm bandwidth) (57). Salt titration data was analyzed as described in Microsoft Excel (57). A minimum of three independent replicates were performed to obtain average K_d(1M)_ and Z_eff_ values for each RNA-protein combination tested, and standard deviations were used to determine statistical significance.

### Mass photometry (MP)

#### a. MP without crosslinking

MP was performed on a Refeyn Two mass photometer run with AcquireMP software. Protein samples were buffer exchanged into glycerol-free buffer (20 mM HEPES pH 7.5, 300 mM NaCl) using Zeba 0.5 mL spin desalting columns (Thermo Scientific) followed by protein concentration determination via A_280_ measurement. Calibration data was collected in MP buffer (35 mM HEPES pH 7.5, 5 mM MgCl_2_, 150 mM NaCl) using soybean β-amylase and thyroglobulin proteins as standards. Np (WT or phosphomimetic mutants) was droplet diluted to a final concentration of 25 nM in MP buffer and movies were recorded for 1 min. Data were analyzed and processed using Refeyn DiscoverMP software.

#### b. MP with crosslinking

WT and 6D Np were buffer exchanged into crosslinking buffer (25 mM HEPES pH 7.5 and 70 mM KCl) using BioRad micro Bio-Spin P-6 gel columns or Amicon Ultra 0.5 mL 10 kDa MWCO diafiltration columns. Concentrations were measured via A_280_ using extinction coefficients listed in Table 1. SARS-CoV-2 RNAs (SL1-4 and SL5) were folded in 50 mM HEPES pH 7.5 at 80°C for 2 min, followed by incubation at 60°C for 2 min, addition of 10 mM MgCl_2_ and incubation at 37°C for 30 min. RNAs were then stored on ice until data collection. To form complexes, the desired Np variant and RNA species were combined to 15 µM final concentration at a 1:1 molar ratio and incubated for 10 min at 25°C. Complexes were then crosslinked at 13 μM via addition of glutaraldehyde (Sigma) at 0.1% (v/v) final concentration. Crosslinking reactions proceeded for 10 min at 25°C and quenched by adding 100 mM Tris-HCl pH 7.5. Crosslinked samples were diluted by 10x in crosslinking buffer immediately before data collection. Samples were droplet diluted to 100 nM final concentration with crosslinking buffer and movies were recorded for 1 min. Data were analyzed and processed using Refeyn DiscoverMP software.

### Limited trypsin digestions

Limited trypsin digests were performed using sequencing grade trypsin (Promega) at a final concentration of 1 ng protease per 1 µg protein target in digest buffer (20 mM HEPES pH 7.5, 150 mM NaCl, 1 mM MgCl_2_, 1 mM DTT, 1% (v/v) glycerol). Each time point used 5 µg total protein and the reaction was quenched by the addition of 2X protein loading buffer (125 mM Tris-HCl pH 6.8, 4% (w/v) SDS, 20% (w/v) glycerol, 5% (v/v) βME) and immediate boiling at 100°C for 2 min. Reactions were analyzed by 15% SDS-PAGE, and visualized using Coomassie Brilliant Blue staining.

### Differential scanning fluorimetry (DSF)

DSF was performed in 20 mM HEPES pH 7.5, 150 mM NaCl using 1.2 µM protein, and “5X” SYPRO Orange dye (Invitrogen) in DMSO (final concentration of 0.1%), as described (58). Three independent replicates (each in technical triplicate) were collected on a QuantStudio 3 qPCR using an iterative heating protocol. Raw fluorescence values from optical filter 3 (excitation: 550 ± 11 nm; emission: 586 ± 10 nm) were exported for fitting. Averaged data were fit to a sigmoidal function in Microsoft Excel to obtain apparent melting temperature (T_ma_) values and the standard deviation was used to determine statistical significance (58).

### Circular dichroism (CD) spectroscopy

CD spectra were collected on a Jasco J-1500 CD spectrometer. Proteins were buffer exchanged twice into CD buffer consisting of 10 mM Tris-H_2_SO_4_ pH 7.5, 150 mM NaF, 10% (v/v) glycerol (59,60). Proteins were diluted to 0.2 mg/mL final concentration and data was collected in 0.1 cm path length quartz cuvettes using three independent aliquots of each protein (61). Spectrometer settings were as follows: 3 accumulations with baseline correction, 100 nm/min scan speed, 1.00 nm bandwidth, 1 nm data pitch, 2 sec digital integration time, and a CD scale of 200 mdeg/0.1 dOD. Baseline was recorded using freshly prepared CD buffer. Spectra were baseline-corrected in the instrument software and exported; averaged spectra are reported with standard deviation, propagated appropriately, at each data point indicated.

### Single molecule optical tweezers (OT)

An 8.1 kb ssDNA molecule was isolated and trapped by an OT system as previously described (62). Each end of the ssDNA was biotinylated for attachment to 1.7 μm streptavidin coated polystyrene beads. One bead was held in a stationary dual beam counter propagating laser trap and the other was held by a glass pipette tip with its position controlled by a piezo electric stage. Moving the beads apart extended the ssDNA, resulting in increased tension along the template, which is measured by displacement of the bead in the trap. The trap is positioned in the middle of a flow initially filled with buffer (10 mM HEPES pH 7.5, 145 mM NaCl, 5 mM NaOH). The system used a feedback loop to adjust ssDNA extension to maintain a constant tension of 10 pN. Buffer containing 100 nM Np was flowed into the sample to allow protein binding. After 100 s of incubation, protein-free buffer was flowed in to remove protein. The experiment was repeated for five biological replicates for each Np variant. In addition to the Np proteins recombinantly expressed and purified from *E. coli*, OT experiments were performed with a commercially available Np expressed and purified from mammalian HEK293 cells (Acro Biosystems). For each experiment, the amplitude of ssDNA compaction was measured both after the initial Np incubation period and after 200 s of Np dissociation in protein-free buffer.

## RESULTS

### Protein and RNA constructs used in this work

Based on published phosphoproteomic data from three independent labs consistently revealing phosphorylation at specific sites within the SRR of the Np linker IDR, six sites were chosen for phosphomimetic Ser/Thr to Asp mutations (Fig. 1) (63–65). The phosphomimetic mutant denoted herein as 3D1 Np contains S176D, S180D, and S184D in the N-terminal portion of the SRR. The 3D2 Np phosphomimetic mutant contains T198D, S202D, and S206D, all in the C-terminal portion of the SRR. A third mutant containing all six of these phosphomimetic mutations is denoted 6D Np.

**Figure 1.**
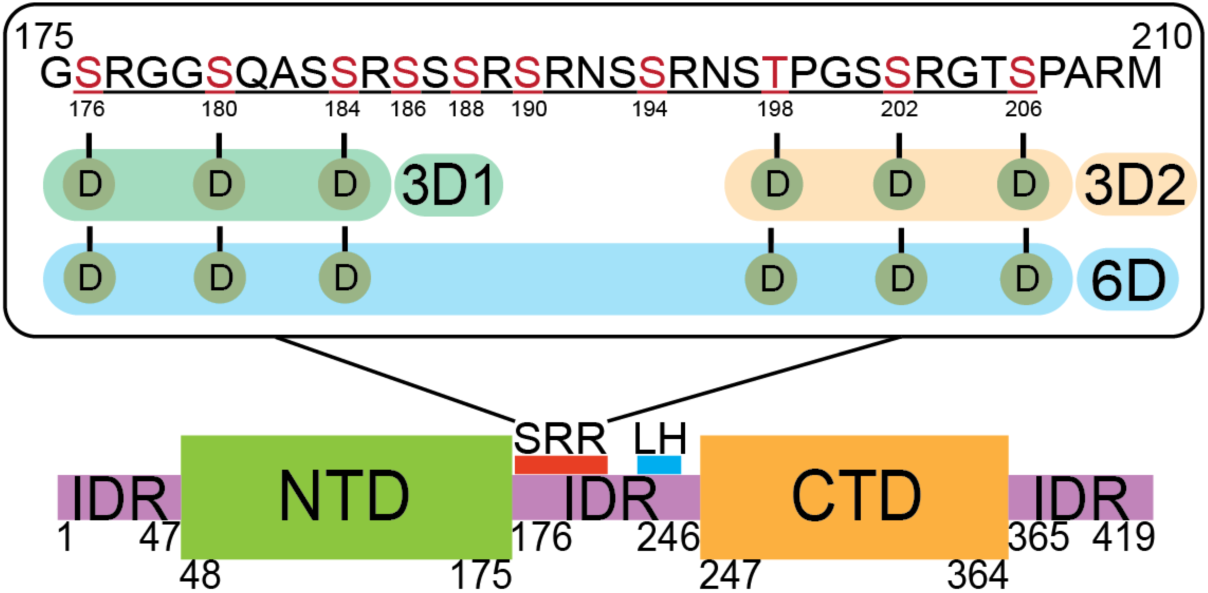
Domain structure of SARS-CoV-2 Np. The red rectangle within the linker IDR is the conserved SR-rich region (SRR) and the blue rectangle highlights the leucine-rich helix (LH); the SRR sequence (ancestral Np) and each of the phosphomimetic constructs used in this work are shown (top): green shading, 3D1; orange shading, 3D2; blue shading, 6D. Red letters indicate all ten SRPK1/GSK3 phosphorylated residues reported by Yaron *et al.* (80).

To study the RNA binding characteristics of these proteins, we used five RNA constructs: three derived from the SARS-CoV-2 5’UTR (Fig. 2A) and two derived from the HIV-1 5’UTR (Fig. 2B). In our salt titration experiments, we also probed HIV-1 full-length Gag for comparison since its binding to the HIV-1 RNA constructs had been characterized with this method previously (43).

**Figure 2.**
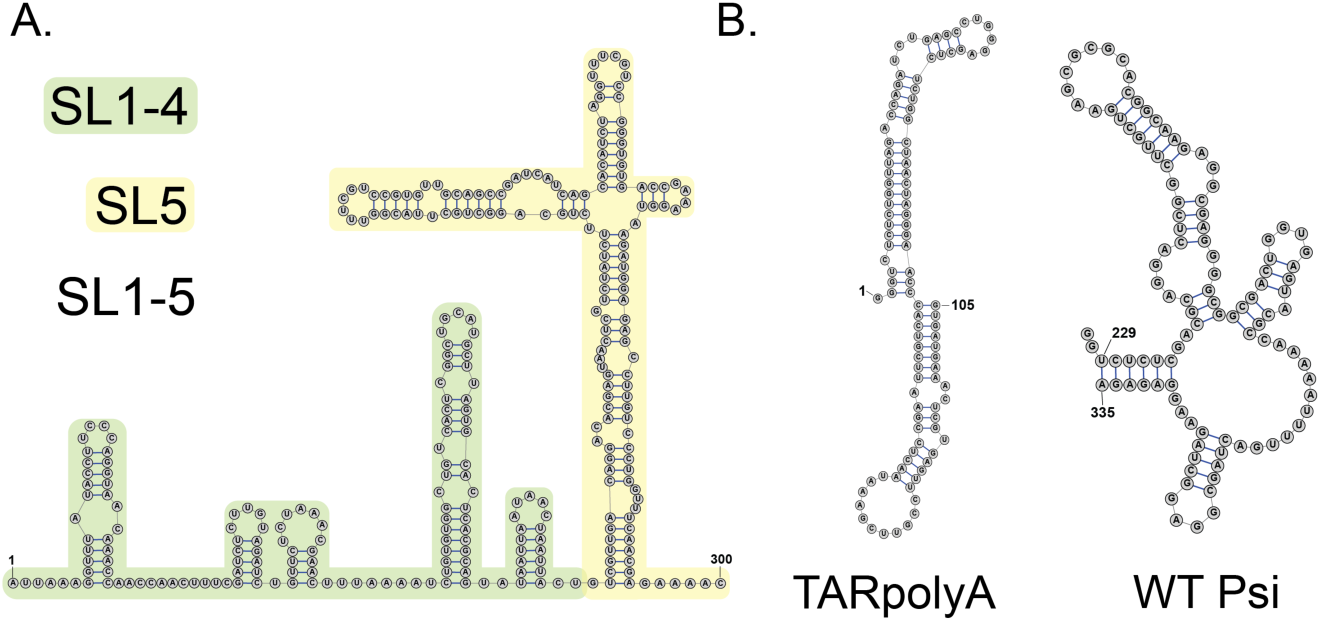
RNA secondary structures of constructs used in this work. A: SARS-CoV-2 genomic RNA 5’UTR (SL1-5, nt 1-300); Green, SL1-4 (nt 1-148); yellow, SL5 (nt 149-300). B: HIV-1 genomic RNA 5’UTR constructs: TARpolyA (nt 1-105); Psi (nt 229-335).

### Effect of phosphomimetic mutations in the central disordered linker on protein structure, dimerization, and stability

To compare WT and phosphomimetic Np structures in solution, we used limited trypsin digestion. Time points were run on denaturing polyacrylamide gels, and after only 30 sec, clear differences between the digestion patterns of WT and 3D2/6D were observed (Fig. 3). Decreased trypsin cleavage of 3D2 and 6D at ∼30 kDa was observed, which is consistent with protection from cleavage, perhaps due to intramolecular salt bridge(s) between phosphorylated sites in the SRR and downstream positive amino acids. At 2 and 5 min, the size of the full-length WT band is smaller than that of 3D1, 3D2, and 6D, implying the non-phosphorylated protein may be more extended and accessible to trypsin. By 10 min, more full-length 6D Np remains relative to the other three, implying that protection increases with an increase in number of negative charges added. Interestingly, the digested band intensity patterns for WT and 3D1 are similar, while the 3D2 and 6D patterns resemble each other (Fig. 3). This suggests that the N-terminal phosphomimetic residues have less of an effect on global structure/protease access than the C-terminal residues.

**Figure 3.**
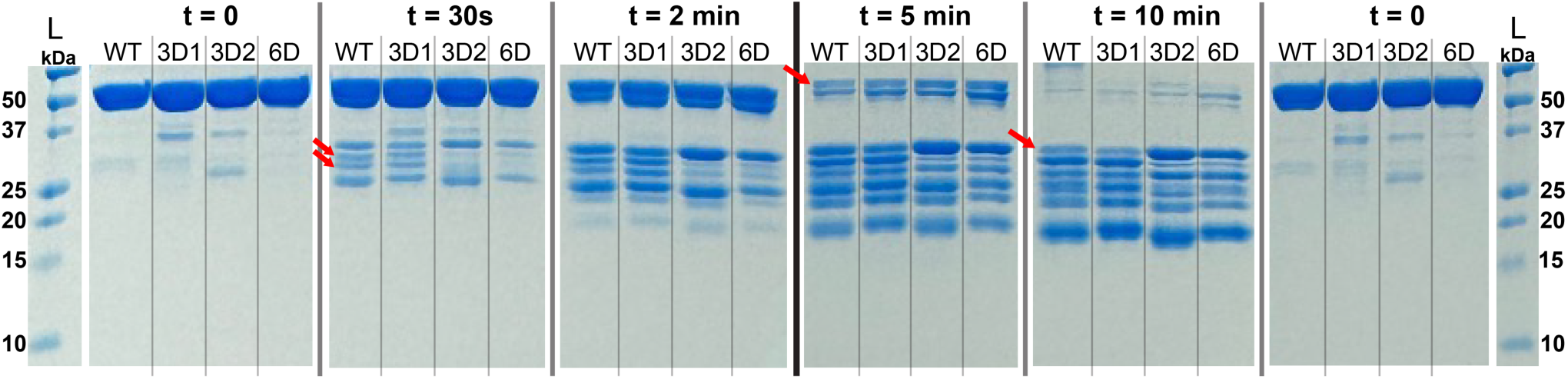
Limited trypsin digests of WT, 3D1, 3D2, and 6D Nps visualized on two 5% / 15% SDS polyacrylamide gels. The ladder and t = 0 loading control are shown at the left and right for both gels, with the central black bar separating them. Red arrows indicate bands that show differences across the tested mutants. Three independent experiments were performed and a single representative replicate is shown.

Based on the altered protease digestion patterns, we hypothesized that thermal stability would be different for the different Np constructs. Apparent melting temperatures were measured for all four proteins using DSF. No statistical difference in T_ma_ was observed between WT, 3D1, 3D2, and 6D Np (Fig. 4). Since DSF measures an increase in the fluorescence of a reporter dye as hydrophobic residues in structured domains are exposed upon protein unfolding, the thermal stability being measured is that of the structured NTD and CTD domains. We conclude that SRR phosphorylation at the residues tested does not affect the overall thermal stability of the structured RNA binding and dimerization domains.

**Figure 4.**
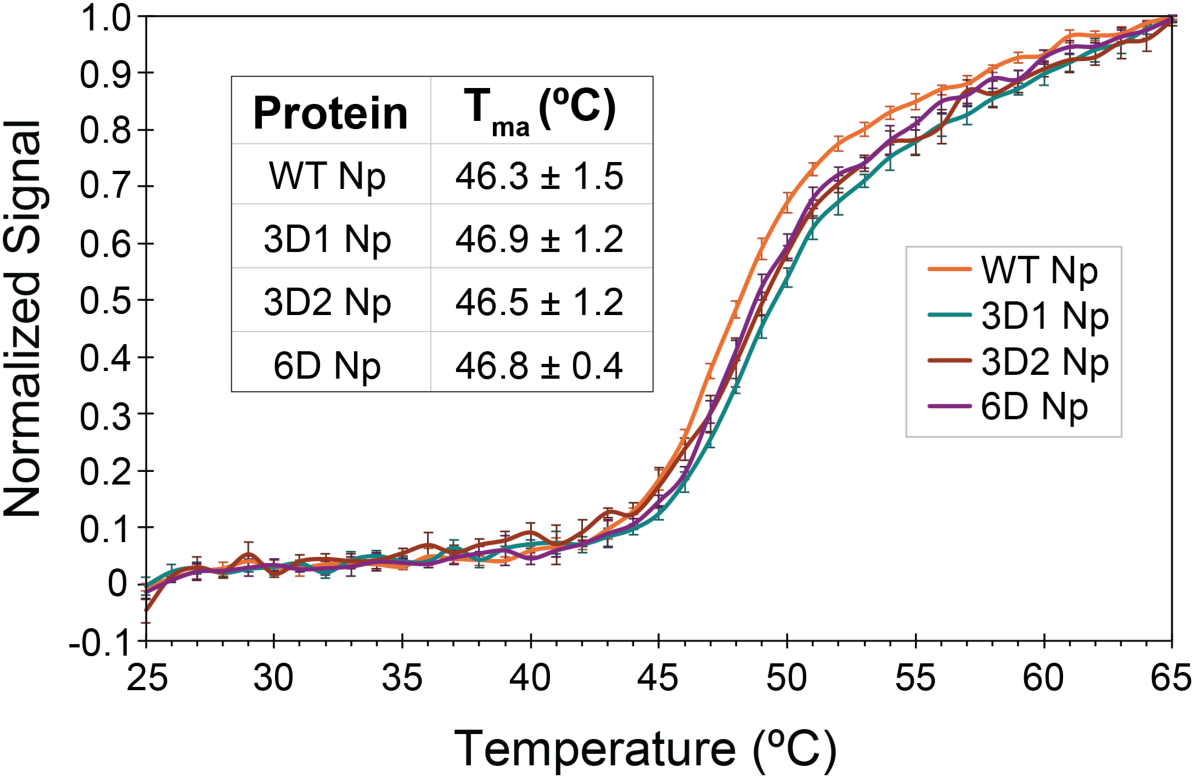
Normalized, averaged DSF curves for WT, 3D1, 3D2, and 6D Nps. Inset shows the calculated average apparent melting temperatures (T_ma_) for each protein with standard deviations indicated. Averages are from three independent replicates.

CD spectroscopy also confirmed little overall change in structure between WT and phosphomimetic constructs (Fig. 5). As previously observed, the CD spectra are consistent with a significant amount of random coil character due to the long IDRs, which constitute 42% of the amino acids (66,67). To better quantify possible differences in secondary structure, we calculated intensity ratios at wavelengths 222 nm/208 nm, indicative of ⍺-helicity (68,69). Although these values do not differ significantly from each other, a decreasing trend is observed with WT having the largest ratio values, followed by 3D1, 3D2, and 6D. The 201 nm peak is indicative of disordered/random coil structure; the difference in the intensity of this peak suggests that the phosphomimetic mutations may cause changes in the disordered regions of the protein (70). Together, these data suggest that phosphorylation of Np does not affect the structured domains but may affect solvent accessibility and potentially local structure in the IDRs.

**Figure 5.**
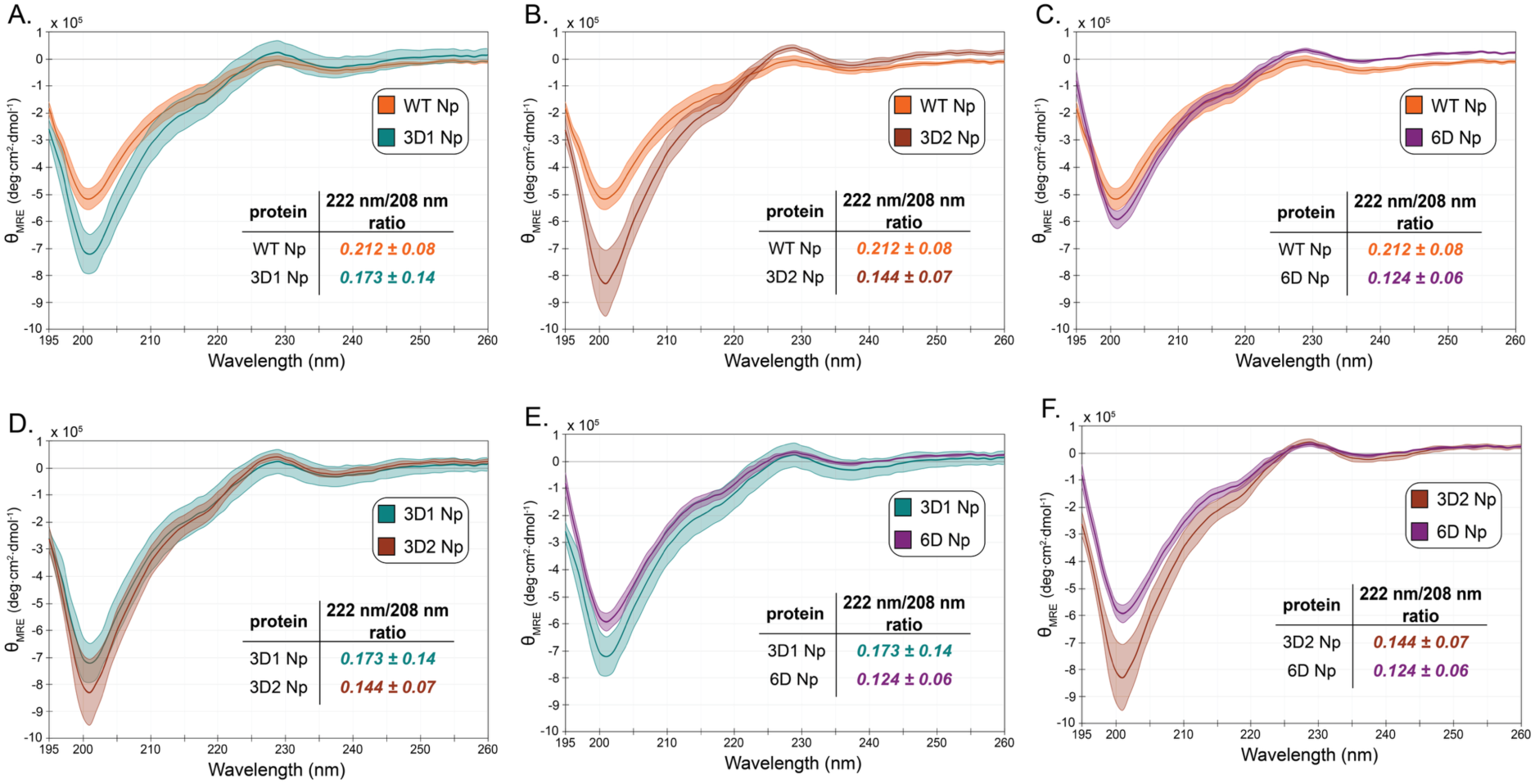
Comparison of CD spectra of WT, 3D1, 3D2, and 6D Nps. Shaded regions show total propagated error for each spectrum after three independent measurements, and the inset table shows the intensity ratio at wavelengths 222 nm and 208 nm with propagated error. Two spectra per plot are shown for ease of comparison among all four proteins. Orange, WT; teal, 3D1; brown, 3D2; purple, 6D. A: WT and 3D1 Np; B: WT and 3D2 Np; C: WT and 6D Np; D: 3D1 and 3D2 Np; E: 3D1 and 6D Np; F: 3D2 and 6D Np.

MP was used to compare each protein’s oligomeric state (Fig. 6). At 25 nM final concentration in the absence of RNA, WT Np is 8% monomer and 91% dimer, whereas 3D1 shows an increase in monomeric species (23%) and reduced dimerization (60%). Interestingly, 3D2 Np closely resembles WT, with 5% monomer and 88% dimer. 6D Np has a monomeric population on par with 3D1 (22%) but 3D1 is the only variant to have a minor tetrameric population (3%) at the concentration tested (Fig. 6). Therefore, phosphorylation of the N-terminal region of the SRR appears to decrease dimerization, and additional negative charges further inhibit oligomerization.

**Figure 6.**
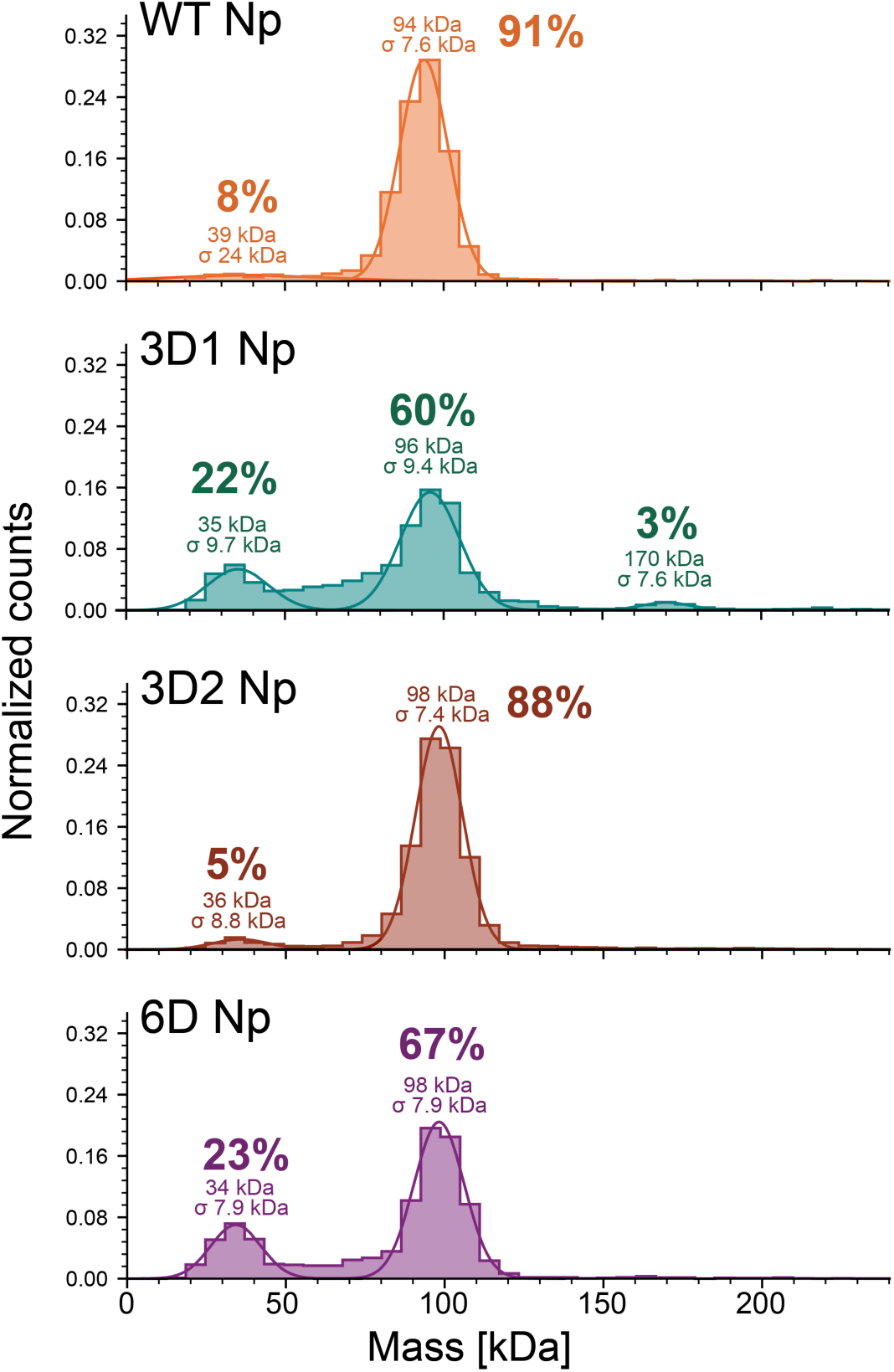
MP histograms of WT, 3D1, 3D2, and 6D Nps at 25 nM final concentration. Histograms shown are combined averages of three measurements. Normalization, averaging, and all other calculations performed using Refeyn DiscoverMP software. Estimated mass of each peak is indicated along with the percent of total counts underneath the Gaussian.

### Effect of Np phosphomimetic mutations on RNA binding

After characterizing the effect of phosphomimetic mutations on Np structure and stability, we next probed differences in RNA binding behavior. Crosslinking-MP was used to determine how Np oligomerization differs in the presence of specific RNAs. Complexes of protein and RNA at a 1:1 molar ratio were formed at 15 µM, crosslinked at 13 μM in the presence of glutaraldehyde, and diluted to 100 nM final concentration prior to data collection.

In the absence of RNA, both WT and 6D Np were exclusively dimeric at 100 nM and the presence of crosslinker stabilized a minor population of tetramer (Fig. 7A and 7B). In the absence of crosslinker, incubation with SL1-4, which has a molecular mass of ∼51 kDa, shows a large RNA only peak in WT but not 6D (Fig. 7C). For both proteins, the major species formed has a mass of 120 kDa with a prominent peak at 200 kDa only observed for 6D Np. Minor higher mass complexes (250-500 kDa range) are also formed with both proteins, even in the absence of cross-linking. The addition of crosslinker resulted in higher order complex formation for both proteins, with complexes up to ∼800 kDa observed for WT Np and up to 1500 kDa with the phosphomimetic variant (Fig. 7D; note difference in X-axis scale). The requirement of crosslinking to observe the large complexes formed between 6D and SL1-4 implies the complexes are reversible in nature. Interestingly, free RNA (∼50 KDa) is only observed in the WT samples (both without and with crosslinking). Given the 1:1 RNA:Np ratio used in our experiments, this suggests that WT Np may bind with a more precise stoichiometry, leaving a certain amount of free unbound RNA. In contrast, in the case of 6D Np, all the RNA participates in the high molecular weight complexes suggesting a broader range of 6D Np/RNA binding stoichiometries.

**Figure 7.**
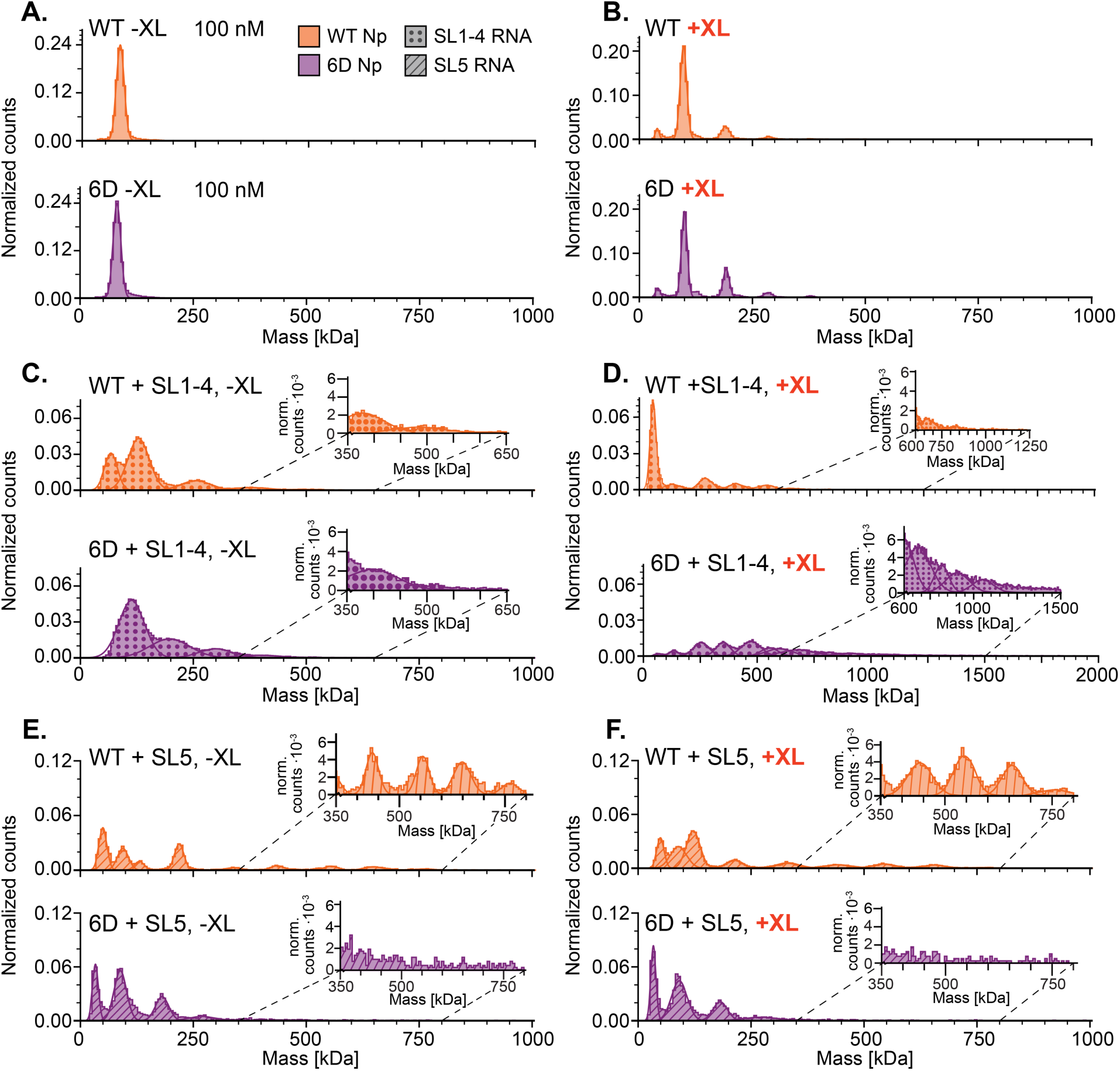
MP analysis of WT Np and 6D Np with (+XL) and without (-XL) crosslinking to SARS-CoV-2 5’UTR RNA constructs. Data were collected at 100 nM final concentrations of each component, and histograms shown are combined averages of three measurements. Normalization and averaging performed using Refeyn DiscoverMP software. A, C, and E: no XL; B, D, and F: with 0.1% glutaraldehyde XL. Note: the x-axis in D is twice that of all others.

In the presence of SL5, which has a similar mass as SL1-4 but is predicted to be more structured (see Fig. 2A), both proteins show an RNA-only peak in the absence of crosslinking but display different binding patterns (Fig. 7E). Minor but discrete complexes with masses up to ∼750 kDa are observed for WT Np. Crosslinking does not significantly affect the size of the complexes observed with SL5 RNA and WT Np but does appear to stabilize the complex at ∼120 kDa and a few of the larger complexes (Fig. 7E versus 7F, top). In the absence of crosslinker, the larger complexes were not observed with WT Np and SL1-4 RNA, which demonstrates reversible binding upon dilution and suggests a weaker affinity for SL1-4 RNA compared to SL5. In contrast to WT, crosslinked 6D Np with SL5 RNA does not show any difference from non-crosslinked (Fig. 7E versus 7F, bottom). In addition, in contrast to the WT protein, without crosslinker, 6D does not show a significant difference in the maximum size of complexes formed with SL1-4 or SL5. However, compared to SL1-4, more of the SL5 RNA remains unbound in the presence of 6D Np without crosslinker; this suggests weaker 6D-SL5 binding (Fig. 7C versus 7E). With crosslinking, 6D Np’s preference for assembly on SL1-4 over SL5 becomes even more clear.

To further characterize the RNA binding characteristics of WT and phosphomimetic Np variants, we performed FA salt-titration binding assays as previously described for HIV-1 Gag binding to the HIV-1 viral RNA 5’UTR elements (43). By fitting the salt titration data to a mathematical model, two values are obtained: K_d(1M)_, which is the K_d_ of the protein-RNA interaction at 1 M Na^+^ when all electrostatic interactions are screened out while the salt-resistant hydrophobic interactions remain, and Z_eff_, which is the number of Na^+^ cations displaced from RNA upon protein binding (57). Therefore, this method can quantify the extent of electrostatic versus hydrophobic binding. In addition to characterizing WT and phosphomimetic Np binding to the three RNA constructs derived from the SARS-CoV-2 RNA 5’UTR (SL1-5, SL1-4, and SL5), binding to two RNA constructs derived from the HIV-1 RNA 5’UTR were examined as non-specific controls: TARpolyA and WT Psi (Fig. 2B). Since Gag binding to the HIV-1 RNAs is well characterized, we also assayed its binding with the three SARS-CoV-2 RNAs for comparison. Fig. 8 summarizes all the binding results in graphical format and average values with standard deviations are found in Supplementary Table S2.

**Figure 8.**
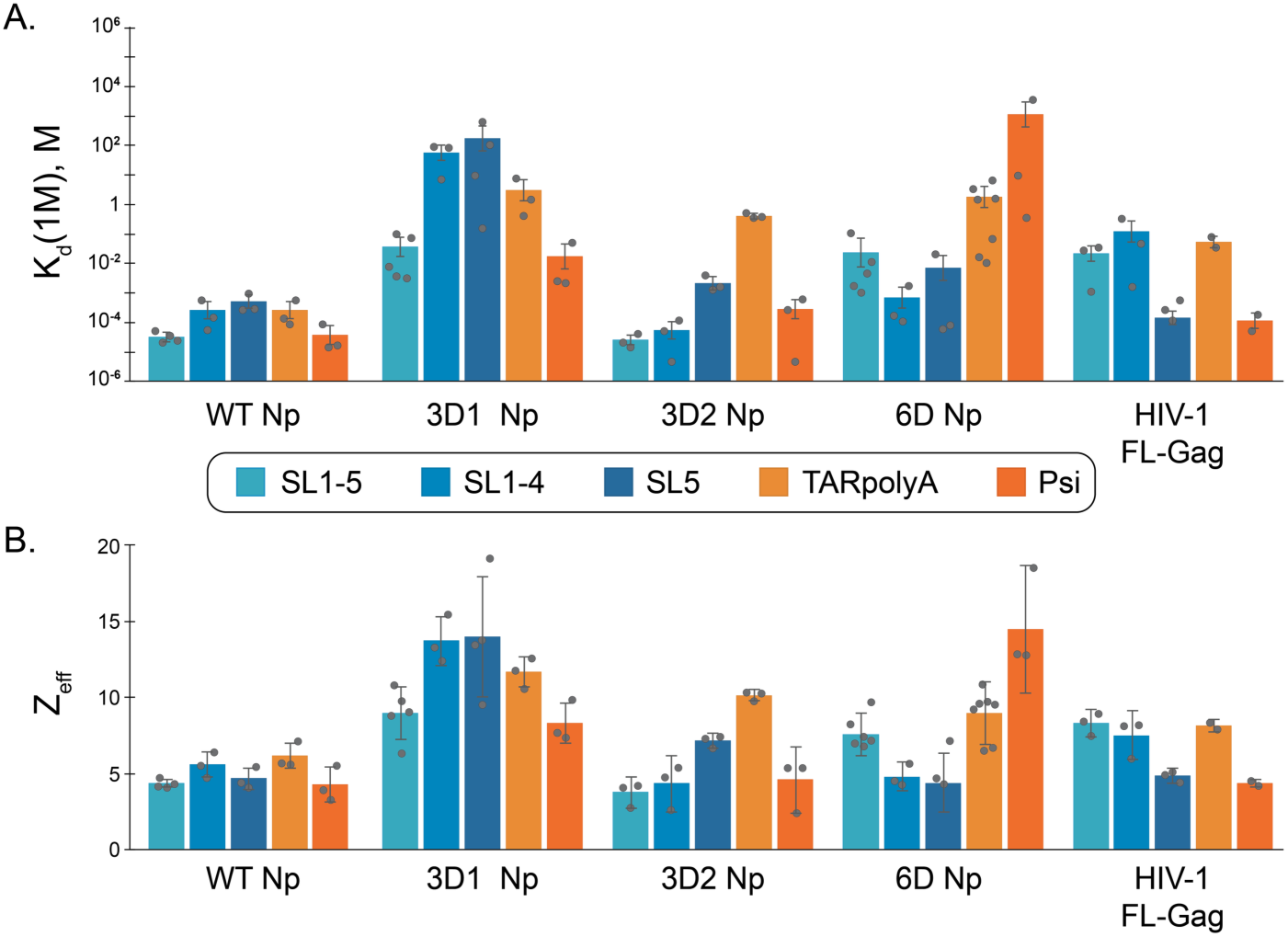
Bar graphs summarizing salt titration parameters obtained for SARS-CoV-2 WT, 3D1, 3D2, and 6D Nps and HIV-1 Gag, color coded by RNA. Individual data points are represented as gray dots. A: Averaged K_d(1M)_ values. B: Averaged Z_eff_ values. All values are the average of at least three independent experiments with bar height representing the mean value.

As expected, K_d(1M)_ and Z_eff_ values are lower for Gag binding to Psi RNA relative to TARpolyA (Fig. 8). This result is consistent with higher specificity of binding to Psi due, in part, to the number of exposed guanosine residues that facilitate specific packaging of the viral genomic RNA (43,71). Interestingly, Gag binding to SARS-CoV-2 SL5 is also characterized by low K_d(1M)_ and Z_eff_ values, possibly due to the presence of several single-stranded (ss) guanosines in loops and bulges, whereas binding to the full-length SL1-5 UTR or to the SL1-4 domain is more similar to TARpolyA binding. Overall, these data are consistent with Gag binding specificity for its own viral packaging signal.

Although non-phosphorylated Np is responsible for viral RNA packaging, whether a specific region of the genome serves as a packaging signal is still unclear (72–75). The K_d(1M)_ and Z_eff_ values for all five RNAs tested with WT Np are relatively low and statistically indistinguishable, suggesting a primarily hydrophobic binding mode (Fig. 8, Table S2).

The phosphomimetic Nps all show quite different behavior, which illustrates that the specific location of phosphorylation in the SR region is important for modulating Np-RNA binding mode. The 3D1 Np construct demonstrated exclusively electrostatic binding behavior, and showed the *highest* electrostatic interactions with four of the RNAs tested (Z_eff_ = 9-14), with the exception of HIV-1 Psi RNA (Z_eff_ = 8.3), where the 6D variant had a higher Z_eff_ of 14.5 (Table S2). For comparison, the highest Z_eff_ value calculated for WT Np was 6.2 with HIV-1 TARpolyA RNA. Conversely, 3D2 Np shows mostly hydrophobic RNA binding to three of the RNAs tested (SL1-5, SL1-4, and Psi) with SL5 parameters suggesting both electrostatic and hydrophobic contributions to binding, and HIV-1 TARpolyA values supporting a primarily electrostatic binding mode.

Interestingly, 6D Np was the only variant tested that differentiated between SARS-CoV-2 5’UTR RNAs and HIV-1 5’UTR RNAs. Whereas WT Np bound to all RNAs tested similarly, 6D bound significantly more hydrophobically and less electrostatically to all three SARS-CoV-2 RNAs while its binding to both HIV-1 RNAs was purely electrostatic. Although 6D K_d(1M)_ values indicate greater specificity of binding to SARS-CoV-2 RNAs than to HIV-1 RNAs, 6D Np’s affinity for SL1-5 is significantly lower than that of WT Np (Fig. 8 and Table S2). Even though the overall positive charge of the phosphomimetic proteins is reduced relative to WT Np, their RNA binding is generally more electrostatic. Taken together, these data suggest that phosphorylation throughout the SRR facilitates Np differentiation of “self” from “non-self” RNA.

Optical tweezers can be used to manipulate and monitor the conformation of a single nucleic acid molecule in real time (Fig. 9A). We have previously used this method to probe nucleic acid binding by full-length SARS-CoV-2 Np and truncation variants. We reported that Np compacted a ssDNA template in two distinct steps and formed stable condensed structures that resisted protein dissociation over tens of minutes (62). In contrast, a truncation construct of Np including only the N-terminal disordered region and NTD significantly decreased Np compaction function and allowed for full protein dissociation within ∼40s.

**Figure 9:**
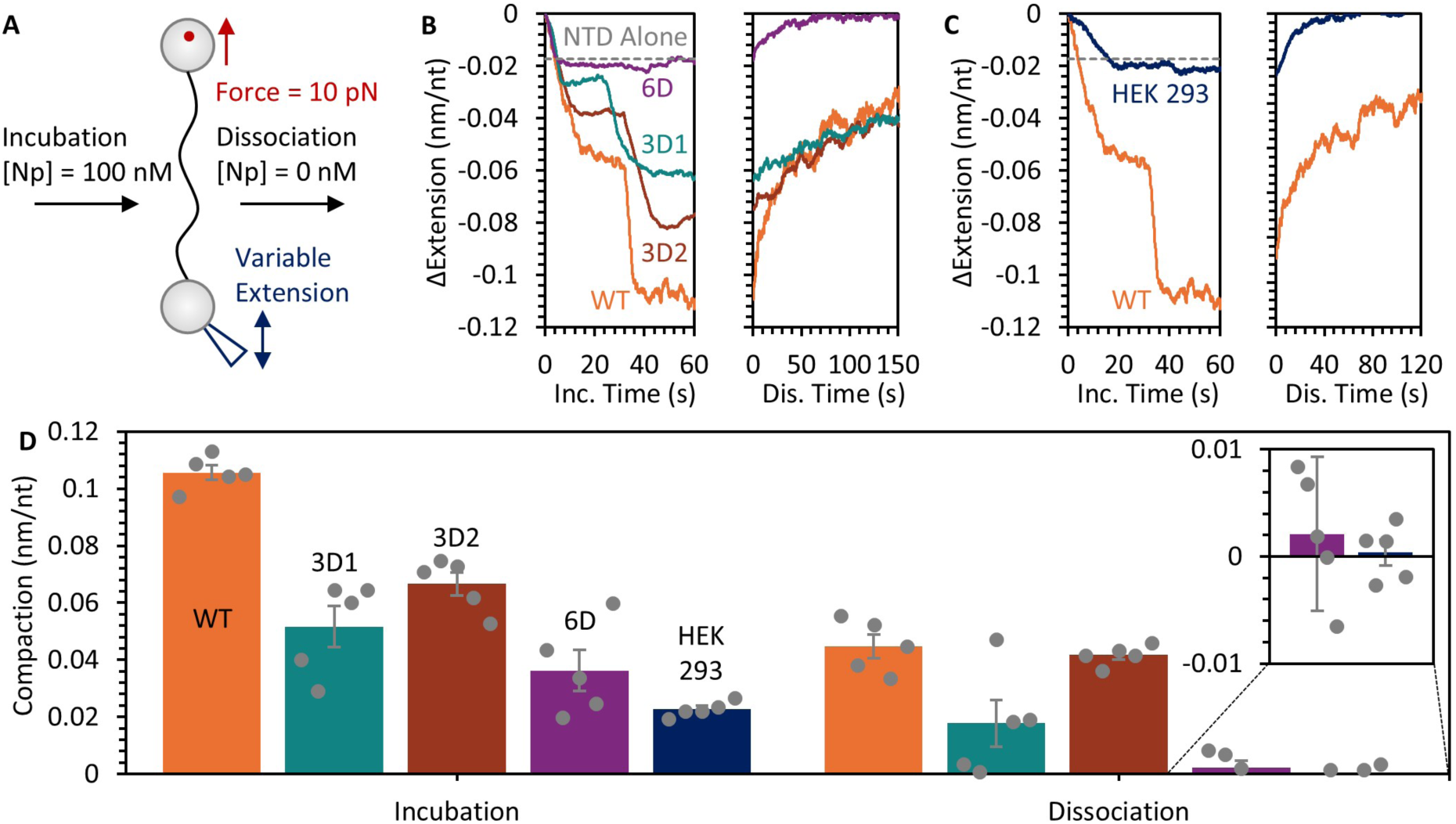
Optical tweezer measurement of WT and phosphomimetic Np ssDNA condensation function. (A) Scheme of the experimental set-up showing a ssDNA molecule tethered between two beads. Free Np is added to the system and incubated for 100 sec. Flowing a protein-free buffer allows dissociation to occur while the extension is adjusted to maintain a constant tension of 10 pN. (B) Extension of the ssDNA molecule monitored over time during incubation in 100 nM Np-containing buffer (left) followed by monitoring dissociation as a function of time in protein free buffer (right). Results are color-coded as follows: WT (orange), 3D1 (teal), 3D2 (brown), and 6D (purple) Np. Previous results with the Np NTD alone (gray dashed line) are shown for comparison (62). (C) Compaction function measured as described in panel B for HEK293T-expressed and purified Np (dark blue). Results for *E. coli*-expressed WT Np are shown for comparison. (D) Summary graph showing the average compaction measured at the end of Np incubation (left) and dissociation (right) based on five biological replicates for each condition with error bars indicating standard error of the mean. Color-coding of the bars is the same as in panels B and C.

Here, we performed OT experiments with WT, 3D1, 3D2, and 6D Np. A single ssDNA molecule was first incubated in a buffer containing 100 nM of each protein for 100 sec, before transferring to a protein-free buffer for 200 sec to allow for protein dissociation. Both 3D1 and 3D2 Np displayed two compaction steps, similar to WT Np, but with reduced amplitude (Fig. 9B). Additionally, the DNA remained in a compact state after removal of free protein, indicating incomplete protein dissociation. 6D Np only produced a single, small compaction step during incubation, and all bound protein readily dissociated. Interestingly, similarly weak compaction capability was also observed for Np expressed and purified from a human cell line (HEK293, Fig. 9C). This Np has the same amino acid sequence as the WT Np expressed in *E. coli*, but functioned more similarly to the 6D Np phosphomimic variant, presumably due to the presence of post-translational modifications including phosphorylation. A bar graph in (Fig. 9D) summarizes the net extension change of the ssDNA following the 100 sec incubation time with each Np variant, as well as the final extension change after 200 sec of protein dissociation. All phosphomimetic Np constructs showed significantly reduced compaction function compared to WT Np and dissociated more readily.

## DISCUSSION

In addition to its well-established role in packaging viral genomic RNA, the multifunctional SARS-CoV-2 Np is also thought to contribute to viral replication and transcription within the RTC. Phosphorylation of its central linker domain has been proposed to regulate the switch between these distinct functions (27,28). To gain mechanistic insights into this functional switch, we characterized biochemical and biophysical properties of non-phosphorylated (WT) and phosphomimetic Np variants. We showed that phosphorylation has only subtle effects on protein structure. The reduced trypsin digestion observed for 3D2 and 6D Np may be the result of intramolecular salt bridge formation upon phosphorylation (Fig. 3). These effects do not result in altered thermostability of the two structured domains, nor do they result in global structural differences as determined by CD spectroscopy (Fig. 4 and Fig. 5).

The Np CTD mediates protein dimerization, and the central linker plays a role in higher-order oligomerization through self-association of the LH downstream of the SRR (Fig. 1) (9,76–79). Stuwe *et al* showed that dimerization of a linker-only peptide was inhibited upon introduction of a single phosphoserine, pS188. Modification at this site primes GSK-3 hyper-phosphorylation at additional sites in the N-terminal region of the SRR, including 184, 180, and 176, which were mutated to Asp in our 3D1 construct (40,80). Whereas pS188 in a linker construct appended to the CTD failed to inhibit the LH self-association, the interaction was eliminated upon GSK-3 hyper-phosphorylation (40). Our MP data showed a slight decrease in dimer populations in the 3D1 and 6D phosphomimetic mutants, whereas 3D2 has a similar dimerization propensity as WT Np (Fig. 6). Taken together, these data suggest that Np oligomerization in the absence of RNA can be tuned by specific phosphorylation events within the N-terminal region of the SRR.

Botova *et al* showed that an Np truncation construct (consisting of the NTD, linker IDR, and CTD) phosphorylated by the specific kinases SRPK-1 and GSK-3 reduced RNA binding. The same construct phosphorylated at non-specific residues by PKA showed RNA binding similar to the non-phosphorylated protein, demonstrating a site-specific effect (79). Stuwe *et al* also showed that modifying a single pS188 and subsequent GSK-3 hyper-phosphorylation attenuated RNA binding of both a linker-only or linker-CTD construct; this was not observed when the full-length Np was phosphorylated or hyper-phosphorylated at these sites (40). These data further support position-specific effects of SRR phosphorylation on Np RNA binding.

Our data also show that phosphorylation does not eliminate binding but significantly modulates the binding mode of full-length Np-RNA interactions in a position-specific manner (Fig. 7 and 8). Non-phosphorylated WT Np binds to all RNAs tested similarly and with a predominantly hydrophobic binding mode. N-terminal phosphomimetic mutations (3D1) reduced hydrophobic interactions with all RNAs tested and switched the binding mode to mostly electrostatic. In contrast, C-terminal phosphomimetic mutations (3D2) minimally impacted hydrophobic interactions. Only 6D Np displayed a statistically significant difference between self (SARS-CoV-2-derived) and non-self (HIV-1-derived) RNA binding parameters, suggesting that phosphorylation across the entire SRR domain is important for more selective binding.

Np:RNA crosslinking MP data also support the hypothesis that phosphorylation modulates Np RNA binding behavior, and together with single-molecule data, provide insights into properties that promote either RNA chaperone or RNA packaging functions. Crosslinked non-phosphorylated WT Np formed complexes with both SL1-4 and SL5, indicating little preference for RNA identity (Fig. 7), as expected for a packaging protein in a system where the viral RNA is extremely long (30 kB) and expressed at high levels (81,82). This contrasts with HIV-1, where the 9.2 kB viral RNA genome is present at far lower levels than host cell mRNAs, requiring a mechanism for selective packaging such as specific Psi RNA packaging signal recognition (see Fig. 8 and Table S1) (83). In contrast, 6D Np preferentially bound and extensively oligomerized on SL1-4 over SL5 (Fig. 7); SL1-4 has more single-stranded character than SL5 and contains the L-TRS sequence required for RdRp synthesis of genomic and subgenomic RNAs, which may contribute to this preference (84). The non-discrete binding of 6D Np on SL1-4 over a broad mass range, especially in the presence of XL, is indicative of heterogeneous complexes with a wide range of stoichiometries, in line with dynamic binding. Taken together, these properties are consistent with chaperone function.

The robust ssDNA compaction and limited dissociation capability of WT Np is also consistent with RNA packaging functionality. As previously mentioned, in the absence of nucleic acid, non-phosphorylated Np dimerizes/oligomerizes more extensively than phosphorylated Np constructs; these self-interactions appear to mask selective RNA binding, which can be observed in 6D Np, in favor of tight, irreversible compaction of nucleic acids. Both 6D Np and HEK293T cell-purified Np display poor ssDNA compaction capability and rapid nucleic acid dissociation. The compaction properties and on/off nucleic acid binding kinetics of SARS-CoV-2 Np binding to dsDNA was also shown to be phosphorylation-dependent (85). The innate immune response cGAS protein was unable to compete off de-phosphorylated Np from a dsDNA substrate, but it could compete off a hyperphosphorylated SARS-CoV-2 Np purified from Sf9 insect cells (85). Phosphorylated or phosphomimetic Np’s reversible nucleic acid binding properties align more with chaperone and strand-transfer function than with genome packaging.

Overall, non-phosphorylated SARS-CoV-2 Np is a promiscuous RNA-binding protein that irreversibly and tightly compacts nucleic acids, and these properties align with the primary function of packaging a 30 kB genome into an ∼100-nm virion (86,87). In contrast, phosphorylation at specific residues alters both the oligomerization propensity and RNA binding behavior of Np in a manner that favors selective and reversible nucleic acid binding and chaperone function. Phosphorylated Np’s proposed role in viral replication requires selectivity, as the recognition of the correct region of the viral genome for the discontinuous transcription jump is essential to maintain the correct reading frame for translation. HIV-1 nucleocapsid (NC) protein, a robust nucleic acid chaperone, shares many of the same traits as phosphorylated Np including hydrophobic binding to specific RNAs, semi-reversible dsDNA compaction, and a more electrostatic binding mode for nonspecific RNA interactions (88,89).

The SARS-CoV-2 Np examined in this study represents the ancestral amino acid sequence. More recent and more transmissible variants harbor mutations within the central IDR linker, which recent studies have shown to alter phosphorylation extent and enhance viral infectivity (77,78,90,91). The mechanistic insights gained here provide a framework for understanding how modulation of Np phosphorylation may shift the balance of its dual roles—RNA packaging and replication/transcription—ultimately influencing disease progression and pathogenicity.

## ACKNOWLEDGEMENTS

We thank Dr. Thomas Magliery for discussion of the CD spectra, which aided in our interpretation of the data, and Dr. Yu-Ci Syu for insightful discussion during the development of the Np denaturing purification protocol.

## AUTHOR CONTRIBUTIONS

Megan S. Sullivan: Conceptualization, data curation, formal analysis, investigation, methodology, validation, visualization, writing – original draft, writing – review & editing

Michael Morse: Conceptualization, data curation, formal analysis, investigation, methodology, validation, visualization, writing – original draft, writing – review & editing

Kaylee Grabarkewitz: Data curation, investigation, methodology, validation, writing – review & editing

Dina Bayachou: Investigation, validation

Ioulia Rouzina: Conceptualization, formal analysis, writing – original draft, writing – review & editing

Mark Williams: Conceptualization, funding acquisition, project administration, resources, supervision, writing – review & editing

Karin Musier-Forsyth: Conceptualization, funding acquisition, project administration, resources, supervision, writing – review & editing

Vicki Wysocki: Funding acquisition, resources

## SUPPLEMENTARY DATA

Supplementary Data are available online.

## CONFLICT OF INTEREST

The authors declare no conflict of interest.

## FUNDING

This work was supported by National Institutes of Health grants R01 AI 153216 (to K.M.-F), T32 GM 144293 (to K.G.), P41 GM128577 instrumentation supplement (to V.W., Refeyn Two^MP^), and RM1 GM149374 (to V.W.), as well as National Science Foundation grant MCB-1817712 (to M.C.W.).

## DATA AVAILABILITY

All data described are contained within the article. Requests to access the datasets should be directed to KM-F.

## Notes

### Competing Interest Statement

The authors have declared no competing interest.

