## Supplemental Data for "Mechanism of SARS-CoV-2 Nucleocapsid Protein Phosphorylation-induced Functional Switch"

Table S1. PCR primers

| Primer name | Sequence | PCR Template |
|---|---|---|
| SL1-4_FWD | 5′-GAAAT*TAATACGACTCACTATAG*GGAAACC-3′ | pIDTSmart-HH-SARS-CoV-2 5′ UTR-FokI (linearized) |
| SL1-4_REV | 5′-mCAGTAATTAGTTATTAATTATACTGCGTGAGTG-3′ | |
| SL5_FWD | 5′-GAAAT*TAATACGACTCACTATAG*GGTCGTTGACAGGACACGAG-3′ | pIDTSmart-HH-SARS-CoV-2 5′ UTR-FokI (linearized) |
| SL5_REV | 5′-mGTTTTCTCGTTGAAACCAGGGAC-3′ | |
| 3D1_FT | 5′-CAGAAGGGGATAGAGGCGGCGATCAAGCCTCTGATCGTTCCTCATCACGTAGTCGCAAC-3′ | pET28α-10xHis-TEV-SARS-CoV-2 WTNp |
| 3D1_FS | 5′-TCATCACGTAGTCGCAAC-3′ | |
| 3D1_RT | 5′-GGAACGATCAGAGGCTTGATCGCCGCCTCTATCCCCTTCTGCGTAGAAGCCTTTTGGC-3′ | |
| 3D1_RS | 5′-CGTAGAAGCCTTTTGGC-3′ | |
| 3D2/6D_FT (make 3D2) | 5′-GAAATTCAGATCCAGGCAGCGATAGGGGAACTGATCCTGCTAGAATGGCTGGCAATGGCG-3′ | pET28α-10xHis-TEV-SARS-CoV-2 WTNp |
| 3D2/6D_FS (make 3D2) | 5′-GAATGGCTGGCAATGGCG-3′ | |
| 3D2/6D_RT (make 3D2) | 5′-TAGCAGGATCAGTTCCCCTATCGCTGCCTGGATCTGAATTTCTTGAACTGTTGCGACTAC-3′ | |
| 3D2/6D_RS (make 3D2) | 5′-TTGAACTGTTGCGACTAC-3′ | |
| 3D2/6D_FT (make 6D) | 5′- GAAATTCAGATCCAGGCAGCGATAGGGGAACTGATCCTGCTAGAATGGCTGGCAATGGCG -3′ | pET28α-10xHis-TEV-SARS-CoV-2 3D1 Np |
| 3D2/6D_FS (make 6D) | 5′-GAATGGCTGGCAATGGCG-3′ | |
| 3D2/6D_RT (make 6D) | 5′-TAGCAGGATCAGTTCCCCTATCGCTGCCTGGATCTGAATTTCTTGAACTGTTGCGACTAC-3′ | |
| 3D2/6D_RS (make 6D) | 5′-TTGAACTGTTGCGACTAC-3′ | |

*Italicized bases: T7 promoter sequence*

Table S2. Salt titration average $K_{d(1M)}$ and $Z_{eff}$ values

| | **HIV-1 Gag** | | **SARS-CoV-2 WT Np** | | **SARS-CoV-2 3xD1 Np** | |
|---|---|---|---|---|---|---|
| | $K_{d(1M)}$ (M) | $Z_{eff}$ | $K_{d(1M)}$ (M) | $Z_{eff}$ | $K_{d(1M)}$ (M) | $Z_{eff}$ |
| SL1-5 | $(2.1 \pm 1.8) \cdot 10^{-2}$ | $8.3 \pm 0.9$ | $(3.6 \pm 1.4) \cdot 10^{-5}$ | $4.4 \pm 0.3$ | $(5.9 \pm 4.4) \cdot 10^{-2}$ | $8.3 \pm 0.9$ |
| SL1-4 | $(1.2 \pm 1.7) \cdot 10^{-1}$ | $7.5 \pm 1.6$ | $(2.6 \pm 2.7) \cdot 10^{-4}$ | $5.6 \pm 0.8$ | $31 \pm 48$ | $13 \pm 2.9$ |
| SL5 | $(1.4 \pm 1.1) \cdot 10^{-4}$ | $4.9 \pm 0.5$ | $(5.0 \pm 3.9) \cdot 10^{-4}$ | $4.7 \pm 0.7$ | $38 \pm 57$ | $12 \pm 2.4$ |
| HIV-1 TARpA | [*]$(2.2 \pm 1) \cdot 10^{-2}$ | [*]$9.1 \pm 0.3$ | $(2.6 \pm 2.5) \cdot 10^{-4}$ | $6.2 \pm 0.9$ | $3.1 \pm 3.9$ | $12 \pm 1.0$ |
| HIV-1 Psi | [*]$(5.2 \pm 1) \cdot 10^{-5}$ | [*]$5.0 \pm 0.2$ | $(3.8 \pm 3.9) \cdot 10^{-5}$ | $4.3 \pm 1.2$ | $(1.8 \pm 2.7) \cdot 10^{-2}$ | $8.3 \pm 1.3$ |

| | **SARS-CoV-2 3xD2 Np** | | **SARS-CoV-2 6xD Np** | |
|---|---|---|---|---|
| | $K_{d(1M)}$ (M) | $Z_{eff}$ | $K_{d(1M)}$ (M) | $Z_{eff}$ |
| SL1-5 | $(2.5 \pm 1.4) \cdot 10^{-5}$ | $3.8 \pm 1.0$ | $(3.1 \pm 5.9) \cdot 10^{-2}$ | $6.9 \pm 2.6$ |
| SL1-4 | $(5.5 \pm 5.4) \cdot 10^{-5}$ | $4.4 \pm 1.8$ | $(6.8 \pm 9.4) \cdot 10^{-4}$ | $4.8 \pm 1.0$ |
| SL5 | $(2.2 \pm 1.4) \cdot 10^{-3}$ | $7.2 \pm 0.5$ | $(6.8 \pm 12) \cdot 10^{-3}$ | $4.4 \pm 1.9$ |
| HIV-1 TARpA | $0.41 \pm 0.1$ | $10 \pm 0.4$ | $17.2 \pm 28$ | $10.1 \pm 1.9$ |
| HIV-1 Psi | $(2.9 \pm 3) \cdot 10^{-4}$ | $6.9 \pm 1.9$ | $50.7 \pm 52$ | $13.7 \pm 1.4$ |

Predominantly hydrophobic interaction

Predominantly electrostatic interaction

Purely electrostatic interaction

[*]Webb, J.A. *et al. RNA*, **2013**. doi: 10.1261/rna.038869.113
